# Extending Protein Language Models to a Viral Genomic Scale Using Biologically Induced Sparse Attention

**DOI:** 10.1101/2025.05.29.656907

**Authors:** Thibaut Dejean, Barbra D. Ferrell, William Harrigan, Zachary D. Schreiber, Rajan Sawhney, K. Eric Wommack, Shawn W. Polson, Mahdi Belcaid

## Abstract

The transformer architecture in deep learning has revolutionized protein sequence analysis. Recent advancements in protein language models have paved the way for significant progress across various domains, including protein function and structure prediction, multiple sequence alignments and mutation effect prediction. A protein language model is commonly trained on individual proteins, ignoring the interdependencies between sequences within a genome. However, biological understanding reveals that protein–protein interactions span entire genomic regions, underscoring the limitations of focusing solely on individual proteins. To address these limitations, we propose a novel approach that extends the context size of transformer models across the entire viral genome. By training on large genomic fragments, our method captures long-range interprotein interactions and encodes protein sequences with integrated information from distant proteins within the same genome, offering substantial benefits in various tasks. Viruses, with their densely packed genomes, minimal intergenic regions, and protein annotation challenges, are ideal candidates for genome-wide learning. We introduce a long-context protein language model, trained on entire viral genomes, leveraging a sparse attention mechanism based on protein–protein interactions. Our semi-supervised approach supports long sequences of up to 61,000 amino acids (aa). Our evaluations demonstrate that the resulting embeddings significantly surpass those generated by single-protein models and outperform alternative large-context architectures that rely on static masking or non-transformer frameworks.

## 1 Introduction

Deep learning architectures have fundamentally changed the way we analyze genomic data. The field has evolved from recurrent neural networks (RNNs) to transformer models, revolutionizing our ability to interpret protein sequences [1, 2]. This advancement has spawned powerful protein language models such as ESM-2, ProtTrans, and ProGen, which excel in capturing the complex relationships between amino acids (aa) within proteins [3, 4, 5]. These models create rich, high-dimensional embeddings—mathematical representations of proteins—that have proven to be remarkably effective for various challenging tasks, including annotation of protein functions [6], multiple sequence alignment [7], and structure prediction [8].

However, these advances come with an important limitation: by focusing on individual proteins, current models miss crucial interactions between distant proteins that shape genome functionality. While proximity-based interactions, such as those within operons, have been extensively studied [9], growing evidence highlights the significance of long-range protein interactions across vast genomic distances [10]. These distant relationships present a compelling parallel to natural language structures, where coherence and meaning often depend on connections between distant elements. This biological reality is particularly evident in evolutionary processes—when one protein changes, its partner proteins often must adapt to maintain their collective function [11]. Current transformer-based models, constrained by limited context windows (e.g., ESM-2 is limited to 1,024 aa, or tokens), cannot simultaneously process multiple long proteins. This technical barrier prevents fully understanding and modeling the complex web of protein interactions across entire genomes.

Recent advances in large language models demonstrate that extended context significantly improves a model’s ability to maintain coherence and capture long-range relationships in text [12]. We hypothesize that analogous approaches could yield more accurate protein embeddings and potentially reveal crucial functional relationships between distantly related proteins missed by current methods. However, extending context in language models presents significant computational challenges, primarily due to the quadratic scaling of attention mechanisms with sequence length [13]. While various solutions have emerged—from sparse attention patterns to efficient architectural modifications—these approaches are fundamentally optimized for text processing and do not account for the unique principles of protein biology, potentially limiting their effectiveness when applied to multiple protein sequences.

To address these limitations, we introduce a novel protein language model architecture capable of processing entire viral genomes. Our model achieves a context window exceeding 60,000 aa on a single commodity GPU. This expanded capacity enables capturing both local protein features and genome-wide interactions in their native genomic context. Notably, this context length covers 98% of all known sequenced viral genomes in the NCBI virus database [14] (Figure 13), representing a significant advancement in studying protein relationships at a genomic scale. Our semi-supervised protein language model employs sparse attention mechanisms guided by known protein–protein interactions. This innovation extends the boundaries of protein sequence modeling and provides new insights into long-range interactions shaping viral genome organization and function. Comprehensive evaluations demonstrate that our viral genome embeddings capture biologically meaningful signals missed by current single-protein models, establishing a novel framework for understanding protein relationships in their native genomic context.

## 2 Related Work

### 2.1 Long Context LLMs

Recent advances in large language models have sparked intense research into efficiently handling extended context. Approaches to increasing context length fall broadly into four popular categories [12]: 1) context length extrapolation, 2) hardware and algorithmic strategies, 3) prompt compression, and 4) attention matrix approximation. Context length extrapolation methods enable models to process sequences longer than those on which they were initially trained by manipulating positional embeddings. Hardware and algorithmic strategies focus on optimizing computation and memory usage through architectural innovations. Prompt compression methods reduce the sequence length by condensing the input information into more compact representations before processing. Attention matrix approximation, in contrast, employs computational and statistical strategies to estimate attention mechanisms without incurring the quadratic scaling costs of standard attention. These approaches are discussed in detail below.

#### 2.1.1 Context Length Extrapolation

A fundamental limitation in extending transformer models to longer sequences lies in Positional Encodings (PEs). While PEs are essential for encoding the sequential order of tokens, their effectiveness diminishes significantly when processing sequences that exceed the training data length. This constraint represents a critical bottleneck when adapting models trained on shorter sequences to handle longer inputs.

Early architectures used fixed sinusoidal positional encodings (SinPE) [13], while newer approaches such as rotational position embeddings (RoPE) [15] introduced trainable embeddings that apply position-based rotations to query and key representations. However, traditional PEs suffer substantial performance degradation when processing sequences that are longer than their training context. For example, models that use RoPE or SinPE exhibit significant increases in perplexity, indicating poorer predictive performance, when used to infer sequences that exceed the lengths encountered during training [16]. To overcome this limitation, context length extension strategies have evolved, enabling efficient processing of long sequences without requiring comprehensive model retraining. These solutions primarily utilize two approaches: extrapolation and interpolation. Extrapolation methods, exemplified by Attention with Linear Biases (ALiBi) [16], introduce unlearned static biases into attention calculations, systematically adjusting attention scores between key–query pairs based on distance. In contrast, interpolation methods generalize learned PEs to longer sequences: RoPE-based interpolation linearly scales position indices to match the original context window [17], whereas more sophisticated methods such as Yet another RoPE extensioN method (YaRN) [18] employ nonuniform frequency interpolation to preserve positional information more effectively.

#### 2.1.2 Hardware and Algorithmic strategies

Training transformer models directly on longer sequences would eliminate the need for positional embedding extrapolation techniques. However, this approach is fundamentally constrained by the quadratic memory complexity inherent to the attention mechanism. Specifically, standard attention mechanisms require memory and computational complexity of *O*(*n*^2^*d* + *nd*^2^) per layer, where *n* represents the sequence length and *d* denotes the hidden dimension size. This quadratic scaling imposes substantial hardware limitations: for example, a Tesla V100 GPU with 16 GB RAM can handle significantly shorter sequences compared to an NVIDIA A100 GPU with 80 GB HBM2e memory, assuming identical model architectures and batch sizes.

To overcome these scaling challenges, researchers have pursued multiple complementary approaches. Distributed computing architectures leverage model parallelism, distributing the memory and computation demands of attention across multiple GPUs. This strategy facilitates the processing of longer sequences by partitioning computations across available hardware resources. Another critical approach involves architectural optimization, particularly by balancing the embedding dimension (*d*) and the maximum achievable sequence length.

Additionally, recent algorithmic innovations have substantially improved the efficiency of attention computation without altering its theoretical complexity. For example, FlashAttention [19] optimizes memory access patterns through fused kernel operations implemented in specialized computing frameworks such as Triton [20]. These practical optimizations reduce memory requirements significantly while preserving computational performance, thus enabling the processing of longer sequences within existing hardware constraints. Together, these hardware advancements, distributed computing strategies, and memory-efficient algorithms establish a robust framework for extending transformer models to process substantially longer sequences.

#### 2.1.3 Prompt Compression

Prompt compression methods aim to reduce input dimensionality by exploiting redundancy within input sequences, ideally preserving semantic content. Techniques employed range from straightforward token-level filtering to sophisticated semantic compression algorithms [21]. Recent compression methods have demonstrated compression ratios of up to 95% without substantially degrading downstream task performance [22]. However, this approach introduces a fundamental limitation in protein sequence analysis: compressed inputs cannot generate detailed per-residue embeddings for every amino acid, as compression inherently reduces sequence representation granularity. Consequently, prompt compression is less suitable for tasks requiring residue-level detail, such as protein structure prediction or mutation effect prediction.

#### 2.1.4 Attention Matrix Approximation

An alternative approach to managing extended contexts involves approximating rather than explicitly computing the full attention matrix. This strategy includes two primary methodologies: context-agnostic and context-aware sparse matrices, each offering distinct trade-offs between computational efficiency and representational capacity.

##### Context-Agnostic Sparse Matrices

Context-agnostic approaches address the quadratic complexity of traditional self-attention through systematic approximation techniques [13]. These methods encompass low-rank matrix decomposition [23], sparsity-induced dimension reduction, and simplified softmax normalization [24]. Among the most successful implementations are architectures leveraging predefined sparse attention patterns, exemplified by Longformer, LongLoRA, and BigBird [25, 26, 27], each employing distinct sparsification strategies (Figure 1). Longformer implements dilated sliding-window attention, strategically positioning sparse attention windows around tokens (Figure 1a). LongLoRA extends context length by introducing shifted attention patterns during fine-tuning (Figure 1b). BigBird employs a hybrid approach combining global attention for select tokens, sliding-window attention, and random attention connections (Figure 1c). While BigBird enables whole-sequence consideration, the stochastic nature of random attention introduces variability in performance, as randomly selected attention positions lack inherent semantic or functional relevance. Although these approaches successfully extend context windows, their predetermined patterns remain independent of sequence content and semantic meaning, presenting inherent limitations. Consequently, context-agnostic designs may inadvertently omit critical interactions or excessively emphasize irrelevant sequence elements. Most notably for biological applications, these methods cannot incorporate domain-specific knowledge—such as experimentally validated protein–protein interactions or genomic relationships—into attention calculations.

**Figure 1.**
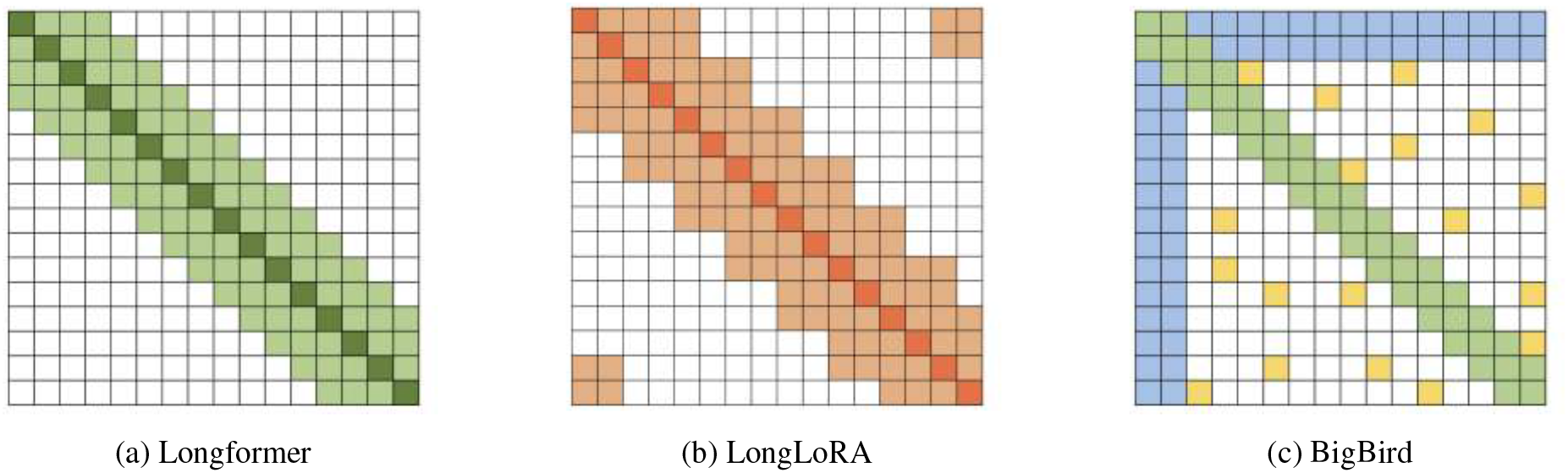
Sparse attention patterns implemented in context-agnostic transformer architectures. **(a)** Longformer’s dilated sliding-window attention pattern, showing a diagonal band of attention with varying intensities of green. **(b)** LongLoRA’s shifted attention pattern during fine-tuning, displayed as orange blocks. **(c)** BigBird’s hybrid approach combining global attention (blue columns), sliding-window attention (green diagonal), and random attention connections (scattered yellow squares). These predetermined sparse attention patterns reduce computational complexity while enabling longer context windows, though they operate independently of sequence content and semantic meaning.

#### Context-Aware Sparse Matrices

In contrast to context-agnostic methods, context-aware compression approaches dynamically adjust their attention patterns based on input content. For example, the Routing Transformer [28] employs K-means clustering on key and query vectors to construct sparse, content-dependent attention matrices. This clustering-based routing enables efficient attention computation between distant sequence elements by restricting attention interactions to intra-cluster relationships. Consequently, memory complexity is effectively reduced from quadratic to linear scaling relative to sequence length, as the attention computation becomes bounded by cluster size rather than the total input sequence length.

However, the exclusive reliance on clustering for routing attention presents significant limitations when analyzing biological sequences. Crucially, clustering-based methods may fail to capture interactions between functionally related but structurally and sequentially distinct elements, such as complementary protein domains that co-evolved despite limited sequence similarity. Indeed, biologically critical interactions frequently occur between proteins exhibiting highly divergent sequences and distinct structural characteristics, challenging the effectiveness of purely clustering-based approaches.

## 2.2 Protein & Genome Language Modeling

### 2.2.1 Protein Language Models

Current popular protein language models (pLMs), including ProtBert [29], ProtTrans [4], and ProGen2 [5], utilize transformer-based architectures with context windows ranging from 512 aa (ProtTrans-XL) to 2,048 aa (ProGen2). These context limitations have largely confined pLMs to analyzing individual proteins in isolation, neglecting their broader genomic context. Among these, the ESM-2 model [3], trained on the UniRef50 database [30] via masked language modeling, has emerged as one of the most widely adopted pLMs. ESM-2 demonstrates impressive performance in protein contact prediction—attributed to its effective attention mechanism—and generates biologically meaningful embeddings useful across diverse downstream applications [6, 7, 8]. However, ESM-2, like its predecessors, focuses exclusively on single-protein sequences with a restricted context length of only 1,024 aa, thereby limiting its ability to capture inter-protein relationships within genomic contexts.

To integrate genomic context into protein representations, the Protein Set Transformer (PST) [31] introduces a graph-based framework wherein proteins, initially embedded via ESM-2, form nodes connected by edges representing genomic proximity. PST propagates contextual information through a multi-head attention mechanism, incorporating learnable positional embeddings that encode protein locations within the genome. By combining local sequence features (ESM-2 embeddings) with genomic relational context (graph representation of proximal interactions), PST refines protein-level embeddings to reflect neighborhood information and generates genome-level embeddings that adaptively weigh protein importance. Despite these advantages, PST relies on precomputed single-protein ESM-2 embeddings and is constrained to modeling interactions between consecutive proteins.

### 2.2.2 Genomic Language Models

Recognizing the inherent limitations of analyzing proteins in isolation has spurred the development of genomic language models (gLMs) [32], which augment ESM-2 embeddings by incorporating context from larger genomic regions. For instance, recent gLMs utilize 30-gene fragments in a masked prediction task, enabling the prediction of protein embeddings by considering the contextual influence of 29 neighboring proteins. This approach generates enriched representations that significantly improve performance across tasks such as protein function prediction and taxonomic classification.

Parallel developments in DNA sequence modeling have yielded architectures capable of processing substantially longer contexts and capturing gene-level dependencies across extensive genomic regions. Prominent examples include Evo [33], DNABert-2 [34], and GenSLM [35], each employing distinct strategies for handling extended sequence lengths. Evo implements a hybrid architecture combining Hyena layers [36] with traditional transformers, initially training on sequences of 8,000 nucleotides and subsequently scaling to 131,000 nucleotides via the StripedHyena architecture [37]. GenSLM adopts a progressive strategy, starting with 2,048-token sequences and extending to 10,000 tokens during fine-tuning using model compression techniques and specialized hardware (Cerebras CS-2). DNABert-2 leverages Attention with Linear Biases (ALiBi) positional encoding, enabling effective inference on sequences significantly longer than its initial 128-token (768-nucleotide) training context without relying on learned positional embeddings. These advances in genomic-scale language modeling represent a meaningful shift from protein-centric analyses, enabling exploration of complex interactions and long-range dependencies across extensive genetic sequences. However, existing gLMs primarily utilize conventional attention mechanisms, highlighting potential opportunities for incorporating biologically informed sparse attention patterns to further enhance their capabilities.

While existing methods have significantly advanced general language modeling and biological sequence analysis, important limitations remain. Current context-extension techniques in language models typically lack biological specificity. Although extrapolation methods such as ALiBi demonstrate effectiveness in general language tasks, they inherently assume consistent structural patterns between short and extended sequences (for instance, local grammatical structures remain stable regardless of sequence length). This assumption does not hold for multi-protein sequences, where relationships between distant proteins can fundamentally differ from the local sequence patterns learned by single-protein models. Additionally, while content-dependent sparse attention mechanisms show promise for managing extended contexts, their potential for explicitly incorporating biological priors into language model architectures remains largely unexplored.

To address these limitations, we introduce a novel genomic-context protein language model specifically designed for viral sequences. Our approach synthesizes recent advances in context-length extension methods with biologically informed sparse attention mechanisms, enabling the effective processing of significantly longer amino acid sequences on commodity hardware while explicitly modeling inter-protein interactions. The following sections detail our methodology and validate its effectiveness across multiple downstream biological tasks.

## 3 Methods

Training protein language models on entire viral proteomes is computationally demanding due to the quadratic scaling of attention mechanisms with sequence length. While existing content-based sparse attention approaches can capture co-evolutionary signals between proteins, these typically utilize static or random masking patterns that neglect biological context. To overcome this limitation, we developed a biologically informed sparse attention method that significantly reduces memory requirements while preserving biological relevance. Our approach computes attention scores exclusively between protein pairs identified as likely interaction partners. Specifically, our methodology involves two main steps:

- Identifying putative Protein–Protein Interactions (PPIs): We utilize attention scores between proteins in shorter viral genome fragments, generated by a fine-tuned ESM-2 protein language model modified with positional extrapolation to handle extended contexts (Figure 2 (a)). Ideally, experimental PPI databases would guide this identification; however, the limited coverage of viral genomes in these databases necessitated this computational strategy.
- Training a novel protein language model on complete viral genomes: We leverage the putative interactions identified in the previous step to train a new, long-context pLM using our biologically focused sparse attention algorithm (Figure 2 (b)). This algorithm strategically restricts attention computations exclusively to protein pairs identified as probable interaction partners.

**Figure 2.**
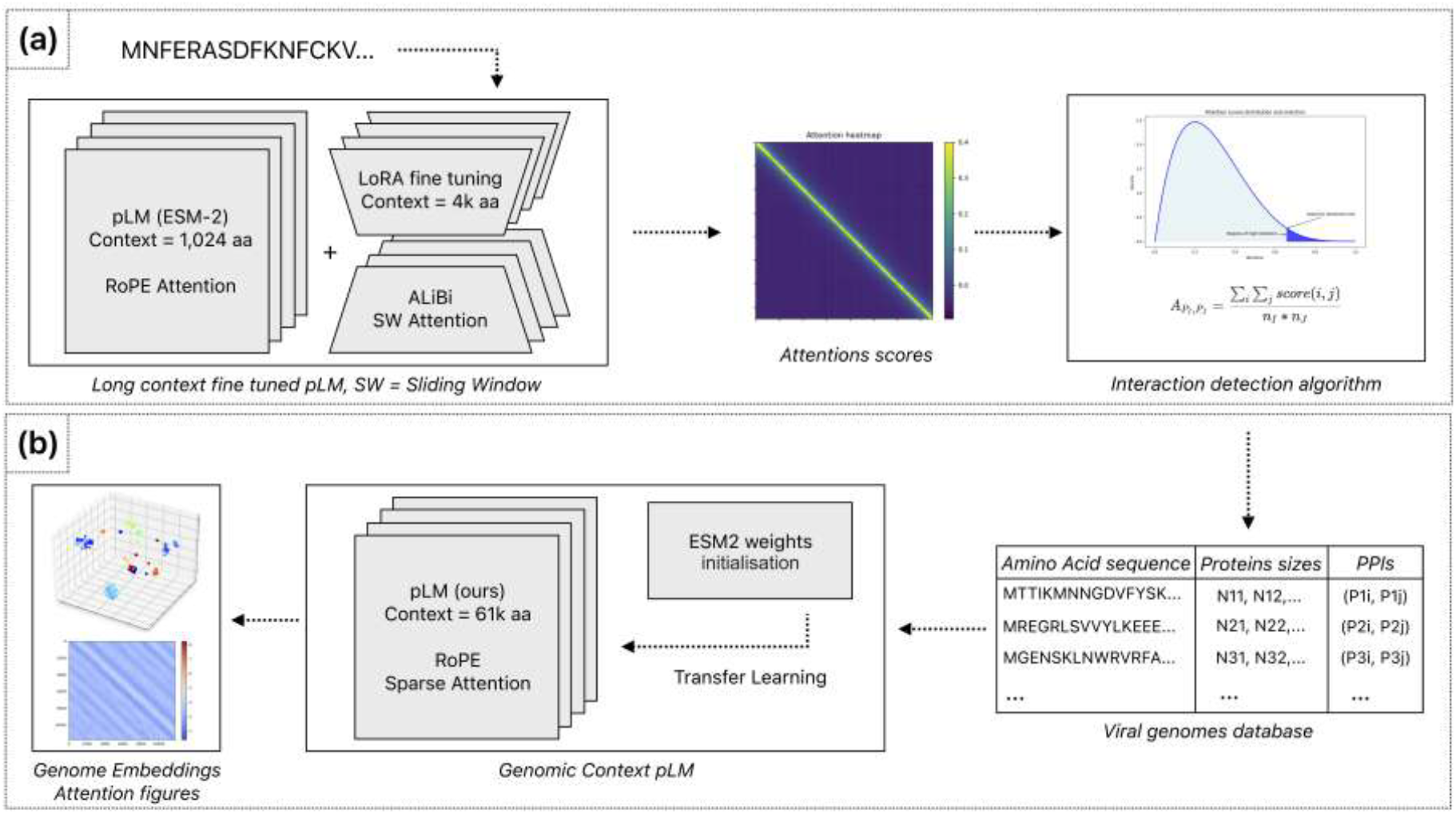
Two-stage approach for training a protein language model on complete viral genomes. **(a)** Identifying putative protein interactions on multi-protein fragments using an extended-context ESM-2 model combining LoRA fine-tuning and ALiBi positional extrapolation. The attention scores produced by this model are subsequently used to inform the protein interaction detection algorithm. **(b)** Training of two long-context protein language models from scratch and initialized from ESM-2 weights using sparse attention patterns guided by predicted protein interactions, enabling efficient whole-genome embedding generation.

**Figure 3.**
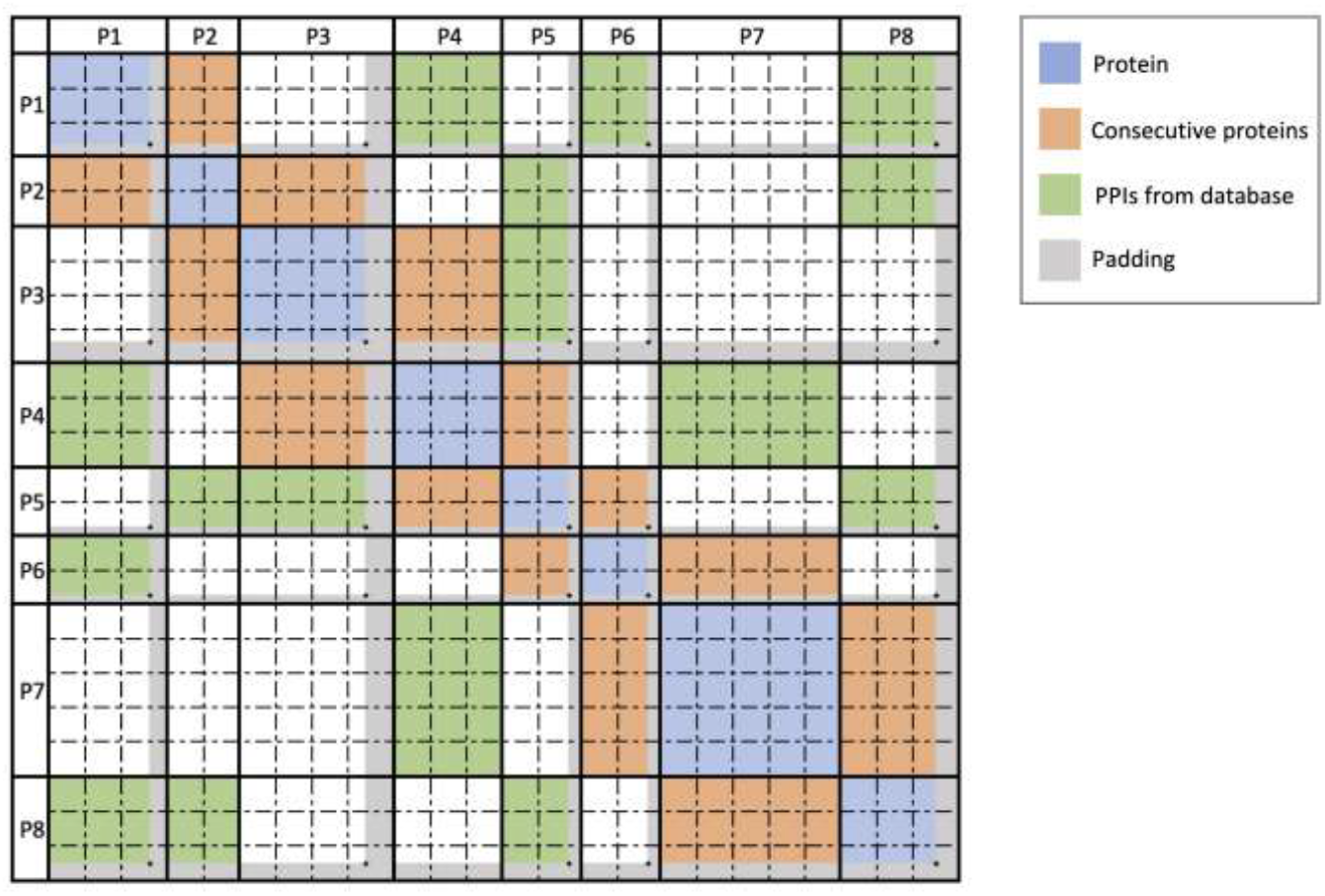
Sparse attention computation architecture. Example attention matrix demonstrating four distinct protein interaction patterns considered during sparse attention computation: intra-protein (self) attention (blue), interactions between consecutive proteins (orange), putative protein–protein interactions identified by our method (section Methods 3.1) and stored in a database (green), and protein pairs not identified as interacting (white), for which attention calculations are omitted. Gray regions indicate padding introduced during fragment construction.

## 3.1 Interactions Inference

PPIs can be inferred from attention scores between proteins in language models, as demonstrated by previous work. For instance, Rives et al. showed that specific attention heads in the ESM-2 model capture distinct protein relationships [38], and Rao et al. demonstrated that self-attention maps encode evolutionary and structural information [39]. Our approach builds upon this concept by analyzing attention patterns simultaneously across multiple proteins within viral genomes to predict potential interactions.

Standard protein language models, such as ESM-2, have a context length limitation of 1,024 aa, restricting analysis to only 3–5 average-length proteins at once. To overcome this, we first extended ESM-2’s context window to 4,000 aa using attention matrix approximation, and subsequently increased it further to 13,000 aa through context length extrapolation (ALiBi). For the initial extension, ESM-2 was fine-tuned on concatenated protein fragments from the NCBI Viral database [14] using an NVIDIA A6000 GPU with 48 GB RAM. Protein fragments were constructed by sequentially concatenating complete proteins in their genomic order until reaching the 4,000 aa threshold, padding as necessary. To enable efficient training on standard GPU hardware, we evaluated multiple sparse attention methods and ultimately selected LoRA due to its proven efficiency in maintaining performance while significantly reducing memory requirements (results not shown).

For each protein fragment analyzed by the fine-tuned model, attention matrices from all encoder layers were extracted. Each attention head (*i*) and layer (*j*)-specific attention matrix *M*_*i,j*_ was normalized and symmetrized using Average Product Correction (APC), resulting in a corrected matrix *N*_*i,j*_ computed as [3]:

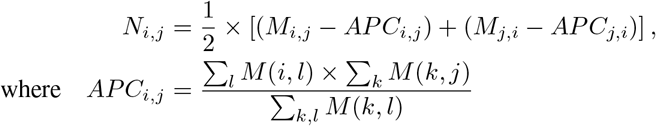

From this normalized attention matrix *N*, we define a set *S* containing the top *t*% highest attention values between amino acid pairs:

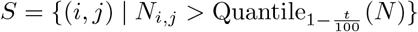

Attention values within the set *S* were subsequently converted to scores via min-max normalization. These normalized scores were then aggregated to compute an attention-based interaction score 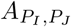 between protein pairs *P*_*I*_ and *P*_*J*_, having lengths *n*_*I*_ and *n*_*J*_, respectively:

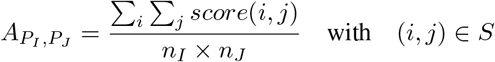

Interaction scores were computed for all protein pairs within the genome. Preliminary analyses indicated that genomically adjacent proteins consistently yielded the highest attention scores (Figure 5), likely reflecting genuine biological interactions. Therefore, for constructing the sparse attention matrix used in subsequent model training, only the top 50 ranked scores from non-consecutive protein pairs were included as interacting proteins, with the remaining pairs masked. This threshold of 50 interactions represents a balance between computational efficiency and interaction coverage appropriate for the viral genome sizes analyzed (Figure 6). The threshold parameter can be adjusted based on genome sequence length and available computational resources.

**Figure 4.**
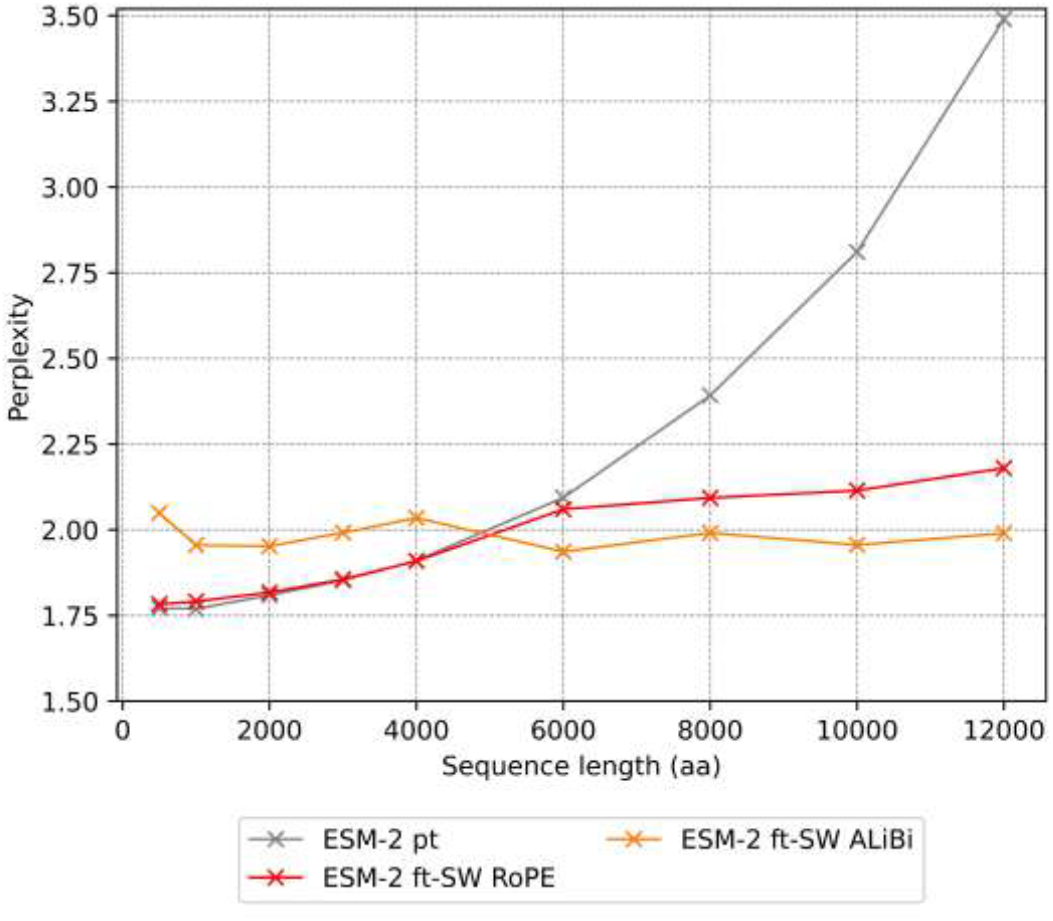
Perplexity evaluation of positional encoding methods for fine-tuning. Performance metrics for ESM-2 variants pretrained (pt) and fine-tuned (ft) using Sliding Window (SW) attention mechanism and two different positional encoding methods(RoPE and ALiBi). Perplexity values represent mean across sequences ranging from 500 to 12,000 amino acids.

**Figure 5.**
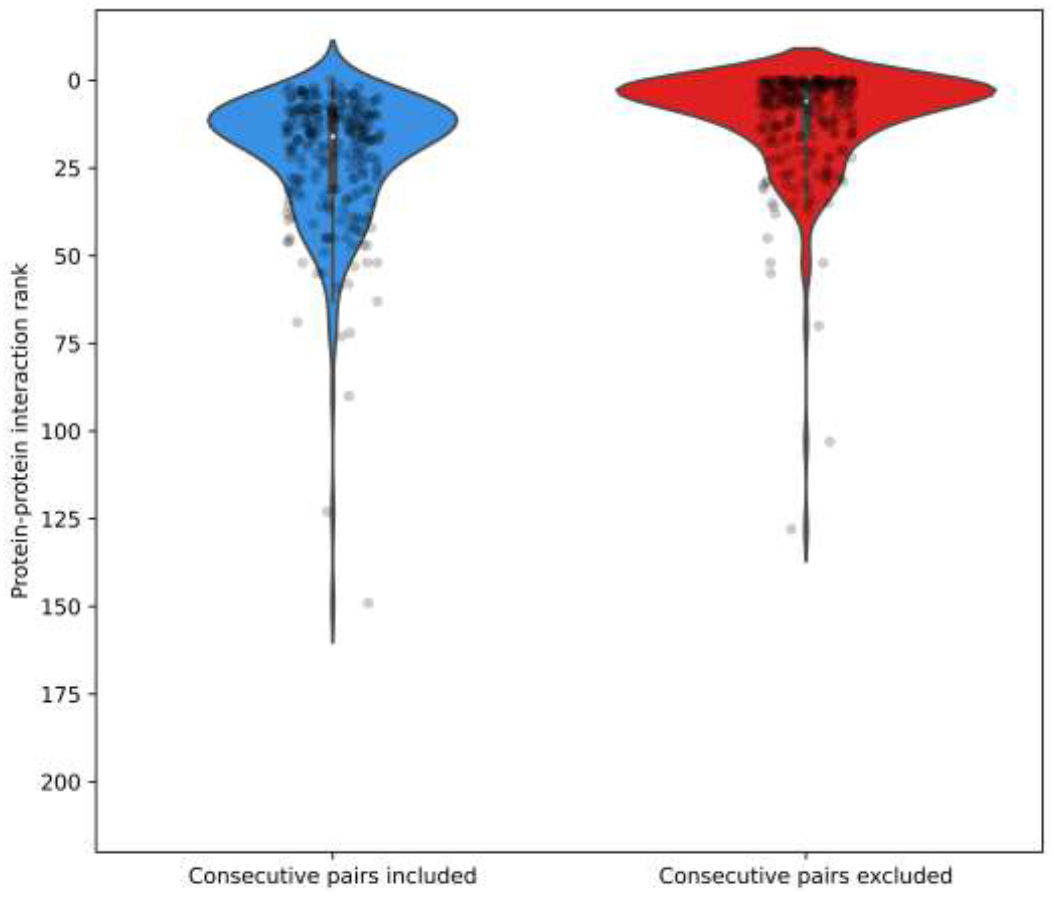
Distribution of interaction pair rankings of DNA polymerase I (PolA), ribonucleotide reductase (RNR), and helicase (HEL) across 83 viral genomes. Violin plots represent the ranking positions of PolA-HEL, PolA-RNR, and RNR-HEL pairs (total n = 249). Left panel: rankings among all protein pairs within fragments. Right panel: rankings after excluding genomically adjacent protein pairs.

**Figure 6.**
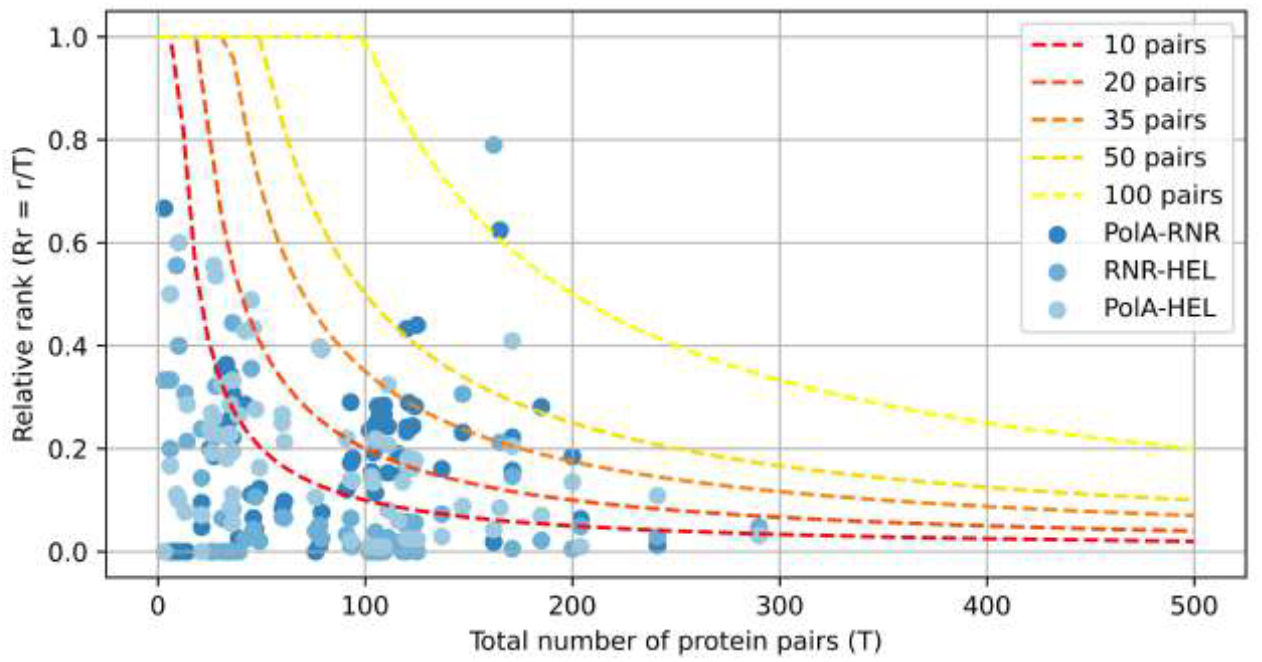
Graph showing selected protein pairs. (PolA-HEL, PolA-RNR, and RNR-HEL) as a function of the total number of genomic pairs and their relative genomic rank (Rr = r/T, ordinate). The dashed curves denote selection thresholds of the top-ranked X pairs. Protein pairs positioned below each curve are retained for subsequent attention calculations.

## 3.2 Sparse Attention Computation

To maintain biological fidelity when computing cross-protein attention, we developed a two-stage fragmentation approach. First, we segment the viral genome precisely at protein boundaries, preserving each protein as a distinct unit. Second, we subdivide each protein into uniform blocks of fixed length *B*, resulting in a block-level representation of the viral genome. We then map the previously identified PPIs onto these blocks, creating fragment-level interaction pairs that guide our sparse attention model. This transformation preserves biologically meaningful relationships while enabling efficient computation across the entire viral genome.

Specifically, given two interacting proteins *P*_1_ and *P*_2_, we transfer these interactions to interactions between their constituent blocks:

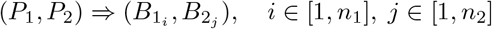

where:

- *P*_1_, *P*_2_ are two interacting proteins from genome *G*.
- 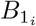 is the *i*-th block of protein *P*_1_, with *n*_1_ total blocks.
- 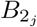 is the *j*-th block of protein *P*_2_, with *n*_2_ total blocks.

For each pair of blocks (*B*_1_, *B*_2_), we compute the block-level attention scores using the scaled dot-product attention formula:

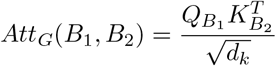

where:

- *B*_1_, *B*_2_ are blocks from genome *G*, spanning from the first amino acid to last amino acids in *P*_1_ and the first amino acid to last amino acids in *P*_2_ respectively
- 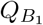 is the query vector subset from genome *G* corresponding to block *B*_1_.
- 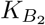 is the key vector subset from genome *G* corresponding to block *B*_2_.
- *d*_*k*_ is the dimensionality of the key vectors.

Efficient computation of sparse attention requires specialized data structures optimized for block operations. We utilize the Block Sparse Row (BSR) tensor format [40], which stores non-zero elements in fixed-size blocks, significantly reducing both memory consumption and computational overhead. Empirical testing identified a block size *B* = 32 as optimal, balancing computational efficiency and minimal padding overhead in our viral protein datasets.

In our implementation, each query–key product 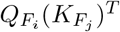 between fragment pairs *B*_*i*_ and *B*_*j*_ produces a 32 × 32 tensor, directly aligning with BSR’s block structure. We subsequently apply softmax normalization followed by multiplication with the corresponding value vectors using optimized Triton [20] kernels tailored for block-sparse operations. Finally, we remove padding tokens to restore original sequence dimensions using precomputed position indices (Algorithm 1).

The training was conducted using the NCBI Virus database (retrieved on 06/2024), comprising 14,436 complete viral genomes totaling 683 million aa. The database represents a comprehensive reference set of viral diversity, ensuring robust generalization across viral families. All genomes were segmented at protein boundaries and padded appropriately to meet sequence length constraints. All training was conducted using NVIDIA A6000 GPUs (48 GB RAM) with block-sparse computations optimized through the Triton library [20].

The two models differ primarily in their initialization and training strategies:

- LV-3C: Randomly initialized and trained from scratch on genomic segments (up to 61,000 amino acids) with an initial learning rate of 3 × 10^*-*4^, using sparse attention informed by inferred protein–protein interactions for 12 epochs.
- LV-5B: Trained leveraging transfer learning from the ESM-2 model (650M parameters) on genomic segments (up to 48,000 amino acids), using biologically-informed sparse attention with an initial learning rate of 3 × 10^*-*5^, also for 12 epochs.

### Algorithm 1

Algorithm for computing context-aware sparse attention, from input tensors through block-wise operations to final output.

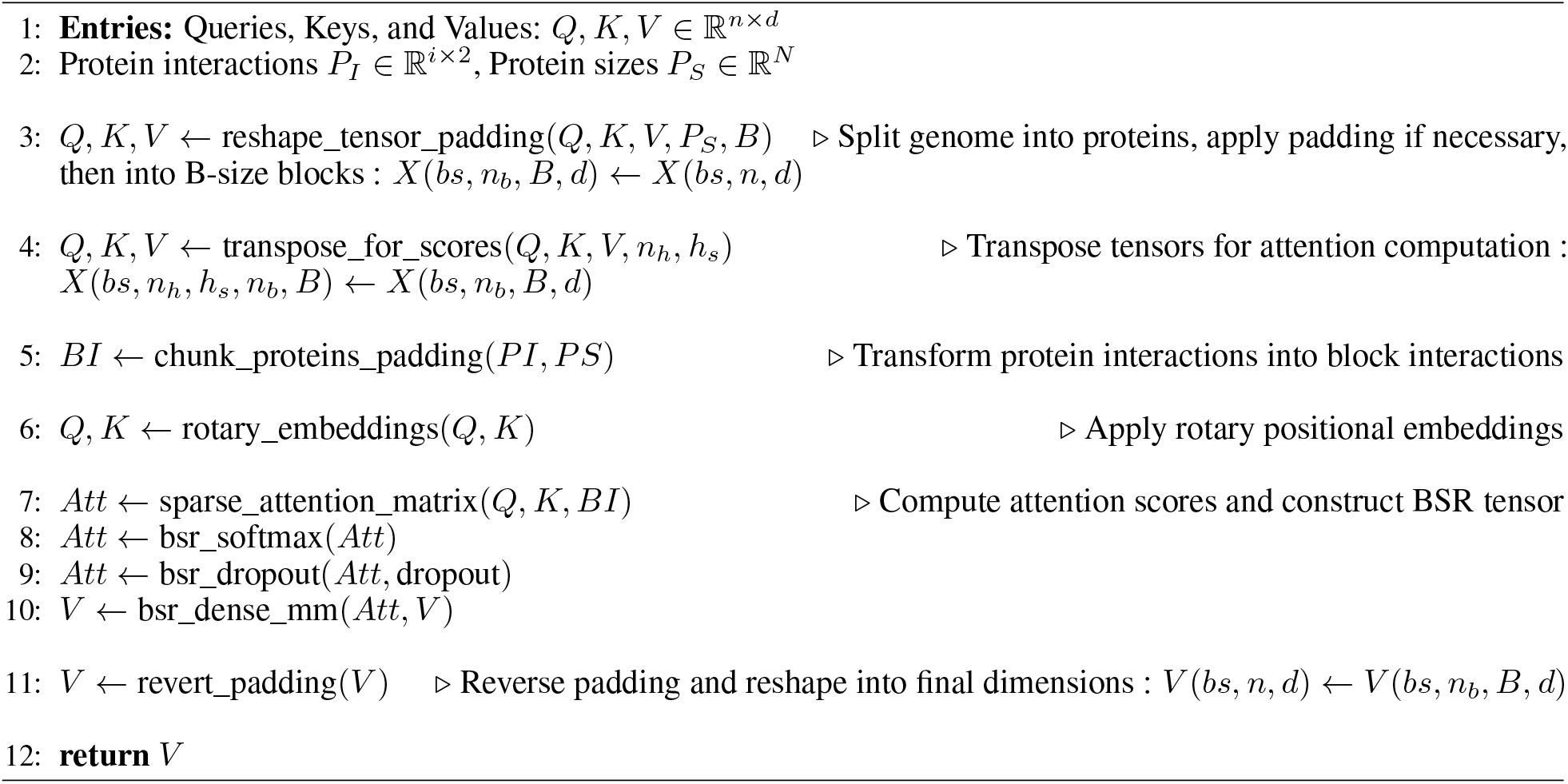

## 3.3 Selecting an optimal fine-tuning strategy

To demonstrate that ESM-2’s context can be marginally extended using existing methods, we evaluated multiple approaches for achieving a context window of 12,000 aa. Specifically, we compared two positional encoding strategies (RoPE and ALiBi) and three sparse attention mechanisms (Longformer, Shifted Sparse Attention, and BigBird). Performance was benchmarked against the baseline ESM-2 650M model using both the perplexity, which quantifies how well the model predicts amino acid sequences (lower is better), and the silhouette scores, which measures how well protein embeddings cluster by structural similarity (higher is better).

To validate these methods on biologically relevant data, we investigated interaction patterns among DNA polymerase I (PolA), ribonucleotide reductase (RNR), and helicase (HEL). These proteins constitute a well-characterized interaction network [41] and display strong evolutionary coupling due to their coordinated roles in DNA replication [42, 43].

## 3.4 Evaluating Global Performance & Embeddings Quality

We evaluated our context-aware sparse attention models (LV-3C and LV-5B) against ESM-2 (pre-trained and ALiBi fine-tuned variants) using perplexity and clustering silhouette scores. Perplexity assessed token prediction accuracy over extended contexts, while the silhouette scores evaluated the quality and information content of protein sequence embeddings. We assessed model performance using viral genome fragments ranging from 500 to 20,000 aa. Although our models support sequences up to 61,000 aa, we limited evaluations to 20,000 aa to match the maximum inference context length of ESM-2 and to accommodate the computational constraints (time and memory) associated with full attention when running on a single GPU (NVIDIA A6000 with 48GB or RAM).

## 3.5 Evaluating Genome Embeddings Quality

We assessed the quality of genome-scale embeddings produced by our models using taxonomic clustering and species classification. To further evaluate embedding utility, we trained a single-layer classifier to identify species from genome embeddings (details provided in Supplementary Material 7).

## 3.6 Evaluation of Embedding Quality for Classification

We evaluated the models’ contextual embeddings by assessing their performance on species clustering and classification tasks, as well as by analyzing the similarity distributions obtained from comparing pairs of embeddings.

- Evaluation of taxonomic information preservation To evaluate how effectively different models preserve taxonomic information, we analyzed clustering patterns of protein embeddings derived from 100 distinct viral species.
- Analysis of species clustering efficiency Next, we trained a single-layer taxonomic classifier designed to predict the species identity of individual proteins directly from their embeddings. To systematically assess classification performance, we varied the number of species classes from 5 up to the 100 species. We measured classifier performance using F1-score averaged over a 10-fold cross-validation scheme.
- Analysis of embedding variance and distribution To further characterize the differences among embedding spaces produced by various models, we randomly selected 5,000 proteins from over 1,000 viral genomes and computed pairwise Euclidean distances between their embeddings. We then compared the resulting distributions from embeddings generated by ESM-2 and our two proposed models, LV-5B and LV-3C.

## 3.7 Evaluating Across-Model Attention

To assess our models’ capacity to capture PPIs within viral genomes, we analyzed attention patterns across the entire network architecture. Attention scores from all heads in each layer were extracted to examine the model’s ability to detect and represent interactions spanning various distances within the genome sequence. We categorized attention interactions into three distinct proximity classes:

- **Short-range interactions**: Attention between amino acids positioned within 100 residues of each other, typically representing intra-protein or intra-domain relationships.
- **Medium-range interactions**: Attention between amino acids separated by 100-500 residues, generally capturing interactions between domains or adjacent proteins.
- **Long-range interactions**: Attention between amino acids separated by more than 500 residues, predominantly representing inter-protein interactions between non-adjacent proteins.

For each category, we calculated the mean attention score across multiple viral genomes, enabling comprehensive analysis of the model’s attention distribution patterns.

## 4 Results

### 4.1 Selecting an Optimal Fine-tuning Strategy

The evaluation of positional encoding strategies showed that ALiBi exhibited superior performance in handling extended sequences. Although RoPE demonstrated robust performance within its original training context, its effectiveness deteriorated significantly beyond 4,000 aa. In contrast, ALiBi maintained consistent perplexity (1.99) across sequence lengths extending up to 12,000 aa (Figure 4).

Among the sparse attention mechanisms tested (i.e., Shifted-Sparse Attention from LongLoRA, Sliding Window Attention from Longformer and Bigbird Attention), Longformer provided the optimal balance between performance (perplexity: 1.99, silhouette score: 0.88) and computational efficiency (training time: 32s/step or 46 hours) (Table 1). Based on these findings, we adopted ALiBi positional encoding combined with Longformer attention, utilizing the LongLoRA method for efficient long-context fine-tuning.

**Table 1.** Performance metrics (perplexity and silhouette score) of ESM-2 pretrained model and several fine-tuned versions using different sparse attention patterns. ‘+full’ means normalization and embedding layers are trainable during low-rank fine-tuning. Metrics are calculated on a pool of genome fragments ranging between 10,000 and 12,000 amino acids.

| | ESM-2 pt | Longformer | Longformer + full | $S^2$-Attn | $S^2$-Attn + full | BigBird |
| --- | --- | --- | --- | --- | --- | --- |
| Training context size | 1,024 | 4,000 | 4,000 | 4,000 | 4,000 | 4,000 |
| Training time (s/step) |  | 32 | 32 | 36 | 36 | 126 |
| Perplexity | 3.17 | 2.18 | **1.99** | 2.22 | 2.05 | 2.02 |
| Silhouette | 0.65 | 0.85 | **0.88** | 0.86 | 0.85 | 0.84 |

We analyzed genomic fragments from 83 viral genomes—each containing complete PolA, RNR, and HEL proteins, which are evolutionarily coupled through their coordinated roles in DNA replication [42, 43, 41]. Consequently, we expect these proteins to exhibit strong inter-protein attention signals in our model. To identify these proteins, we performed BLAST searches using reference sequences for each target protein, retaining only genomes where all three proteins were detected with high confidence. For each selected genome, we extracted the genomic region spanning from the start of the first identified protein to the end of the last one, including all intervening proteins, resulting in fragments of up to 12,000 amino acids in length and averaging 93 proteins per fragment. For every fragment, we computed pairwise interaction scores 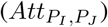 between all protein pairs (*P*_*I*_, *P*_*J*_), and ordered these interactions according to their scores. Subsequently, we assessed the relative rankings of three well-established interaction pairs—PolA-HEL, PolA-RNR, and RNR-HEL—within the complete distribution of pairwise scores (Figure 5).

Figure 5 illustrates that the PolA-HEL, PolA-RNR, and RNR-HEL pairs consistently rank among the highest-scoring interactions identified by our method. Although genomically adjacent protein pairs predominantly occupy the top positions (blue violin plot), removing these adjacent pairs reveals that our proteins of interest still maintain high interaction scores among non-adjacent protein pairs (red violin plot), typically ranking within the top 50 of more than 200 possible pairs. These results corroborate the well-established co-evolutionary relationships among these replication proteins and validate the efficacy of our attention-based interaction detection approach. Additionally, Figure 6 displays the distribution of ranking positions for PolA-HEL, PolA-RNR, and RNR-HEL pairs relative to their genomic rankings.

To identify an appropriate selection threshold, we plotted curves corresponding to thresholds of the top 10, 20, 35, 50, and 100 ranked pairs. Selecting the top 50 non-adjacent pairs retained 97.6% of the known interacting protein pairs (Table 2).

**Table 2.**
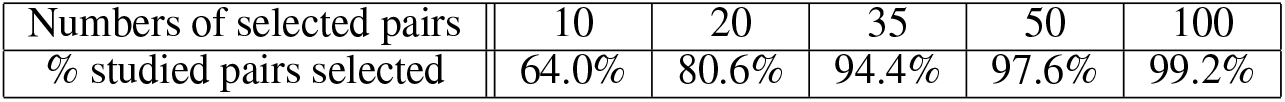
Percentage of studied pairs (between DNA polymerase I (PolA), ribonucleotide reductase (RNR), and helicase (HEL)) selected for different thresholds.

|  |  |  |  |  |  |
| --- | --- | --- | --- | --- | --- |
| Numbers of selected pairs | 10 | 20 | 35 | 50 | 100 |
| % studied pairs selected | 64.0% | 80.6% | 94.4% | 97.6% | 99.2% |

To attain our target context size of 61,000 aa while ensuring computational tractability, we chose the top 50 ranked interactions for the implementation of a content-aware sparse attention mechanism. This threshold effectively captures biologically significant interactions, as demonstrated by our replication protein analysis, while adhering to memory constraints required for processing entire viral genomes.

### 4.2 Evaluating Global Performance & Embeddings Quality

The LV-3C and LV-5B models demonstrated significantly lower perplexity compared to ESM-2 on sequences longer than 3,000 aa (Figure 7a, light blue), maintaining this advantage consistently across all tested lengths (dark blue). In contrast, both the pre-trained ESM-2 (gray) and its RoPE-fine-tuned variant (red) experienced significant perplexity inflation when evaluated beyond their original training context windows. Fine-tuned version of ESM-2 with ALiBi (orange) maintains near-constant perplexity with sequence length, but higher than our models.

**Figure 7.**
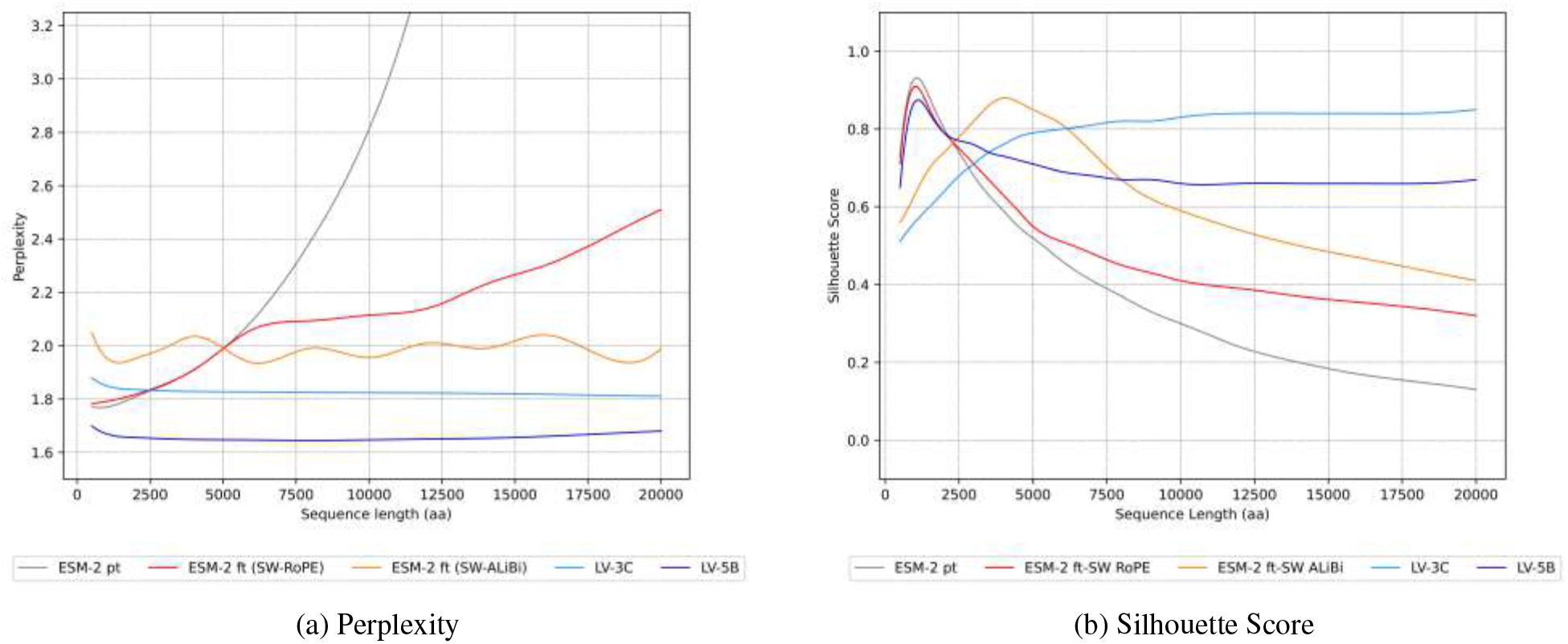
Global performances of our models compared to ESM-2 pt and ESM-2 ft. **(a)** Perplexity evaluation of ESM-2 pretrained, fine-tuned (using RoPE and ALiBi positional encoding strategies), and our models LV-3C & LV-5B. **(b)** Clustering ability evaluation using Silhouette score on fragment embeddings, using sequence similarity as reference with MMSeqs2 clustering. These smoothed curves are obtained from perplexity and Silhouette score measurements for sequences ranging from 500 to 20,000 amino acids.

For clustering quality analysis, we clustered protein sequences and fragments using MMseqs2 [44] at an 80% similarity threshold and computed silhouette scores to quantify embedding coherence. Our results revealed clear differences in clustering performance among the tested models (Figure 7b).

The silhouette curves for ESM-2, its RoPE-fine-tuned variant, and the transfer-initialised LV-5B climb until ≈ 1,020 aa—the maximum length in ESM-2’s pre-training set, signalling improved embedding coherence within this range, but then drop sharply as the encoding remains dependent on the short-range statistics. In contrast, substituting RoPE with ALiBi and continuing training yields a flatter, length-aware attention profile that pushes the decline back to roughly 3,500 aa. Both LV models provided superior embedding coherence and better discrimination of protein sequences compared to baseline approaches. LV-3C (light blue) exhibited robust clustering quality, particularly for longer sequences (silhouette score: 0.83), while LV-5B (dark blue) achieved performance comparable to ESM-2 for shorter sequences and maintained high effectiveness at extended lengths (silhouette score: 0.67). ESM-2 (gray) excelled only within its original training range (1,024 aa), with embedding quality markedly declining for longer, multi-protein sequences. While ALiBi fine-tuning (orange) extended effective clustering performance up to approximately 4,000 aa, embedding quality still deteriorated as sequence lengths increased, despite the positional encoding extrapolation strategy.

### 4.3 Evaluating Genome Embeddings Quality

The quality of the genomic embeddings was assessed by a clustering and a classification task. The aim of the clustering-based evaluation is to identify consistency between embeddings in latent space. This coherence is highlighted here by taxonomic species. Silhouette scores (Figure 8a) indicated strong taxonomic discrimination across all models, with the LV-5B variant consistently outperforming others.

**Figure 8.**
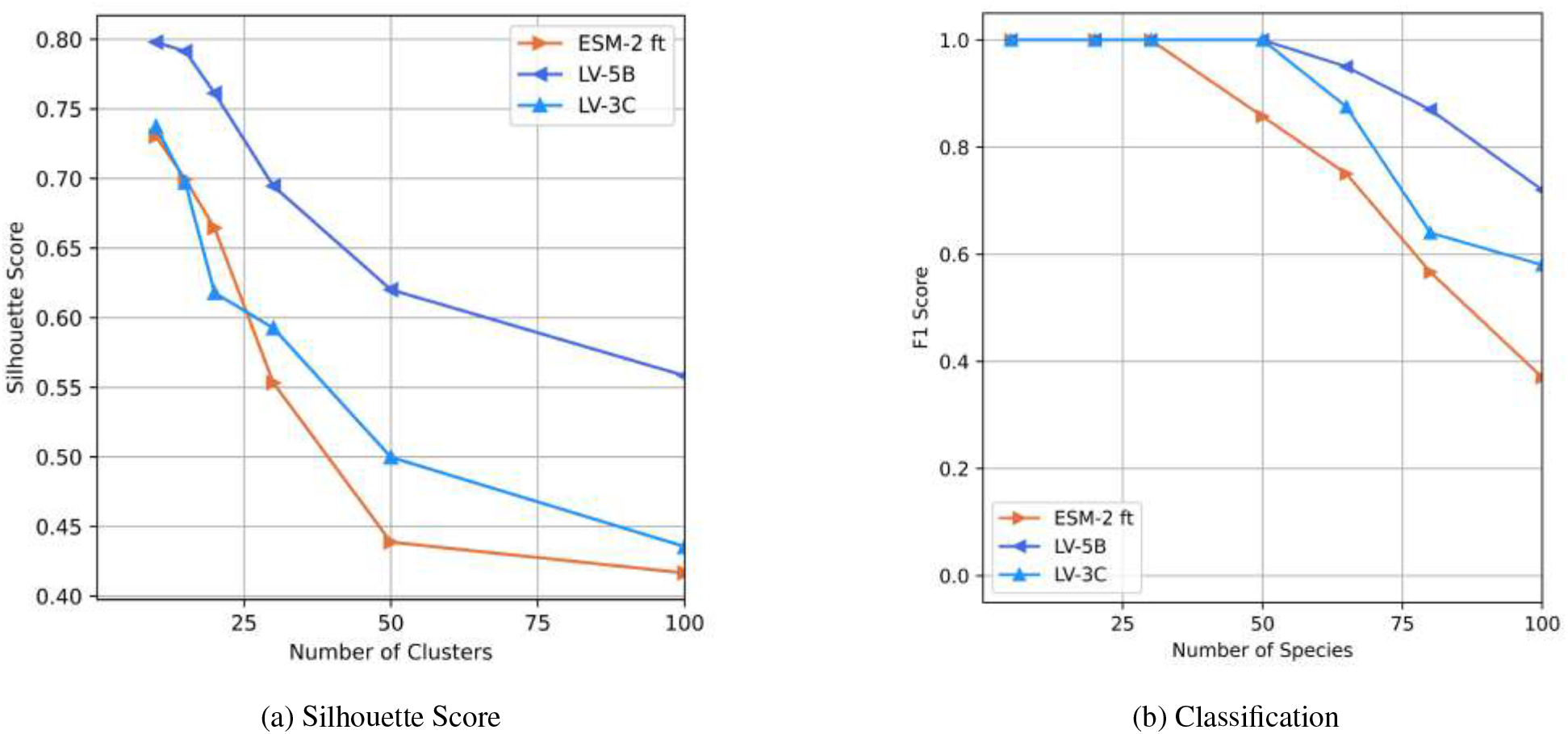
Evaluation of full genome embeddings. **(a)** Silhouette scores comparing genome embeddings generated by LV-3C, LV-5B, and fine-tuned ESM-2 (ALiBi positional encoding), using taxonomic species as cluster labels. **(b)** F1-scores for a single-layer classifier predicting species identity based on genome embeddings from LV-3C, LV-5B, and fine-tuned ESM-2 (ALiBi positional encoding).

The performance of the species classification of genomes evaluated by F1-scores (Figure 8b) revealed a notable trend: as the number of species increased—and consequently, task complexity—embeddings generated using context-aware sparse attention demonstrated superior classification accuracy relative to genome embeddings from fine-tuned ESM-2. This performance improvement at higher taxonomic resolution suggests our embeddings effectively capture nuanced genomic features, enhancing discriminative power at larger phylogenetic scales. These properties hold promising implications for phylogenetic and metagenomic applications.

### 4.4 Evaluation of Embedding Quality for Classification

#### 4.4.1 Evaluation of Taxonomic Information Preservation

Given that protein embeddings inherently cluster primarily by protein type rather than species, silhouette scores across all models were generally low (slightly negative). Despite this challenge, embeddings generated by the LV-3C model exhibited better species-specific clustering relative to those produced by ESM-2 and LV-5B (Figure 9a). For comparison, we included embeddings from the Protein Set Transformer (PST), a recent genome-level protein model, as an additional baseline. PST achieved the lowest silhouette scores, indicating limited capability in capturing viral species-level taxonomic relationships.

**Figure 9.**
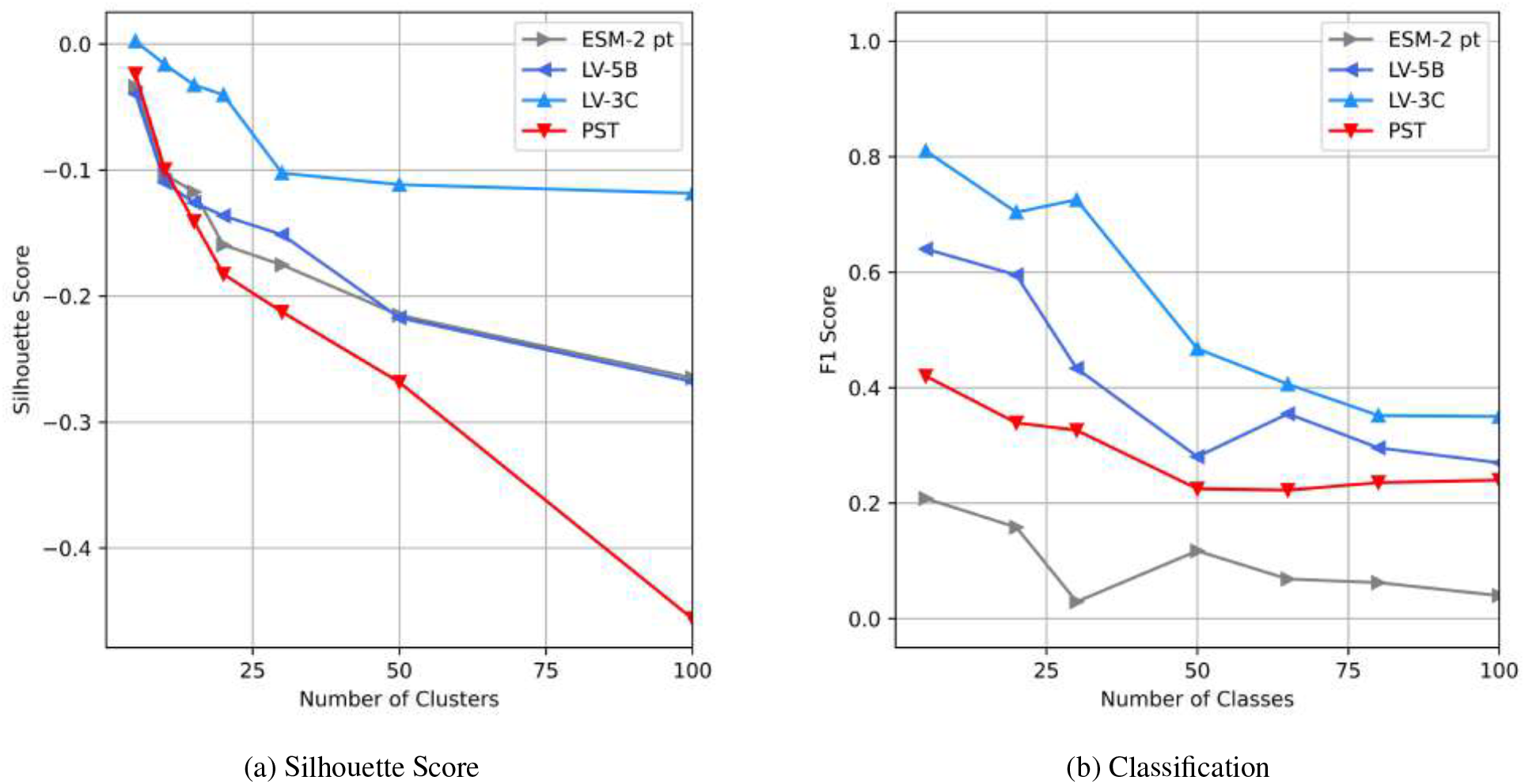
Evaluation of contextualized protein embeddings. **(a)** Silhouette scores comparing protein embeddings generated by LV-3C, LV-5B, Protein Set Transformer (PST), and pretrained ESM-2, using taxonomic species as cluster labels. **(b)** F1-scores for a single-layer classifier predicting species identity based on genome embeddings from LV-3C, LV-5B, PST, and fine-tuned ESM-2 (ALiBi positional encoding).

#### 4.4.2 Analysis of Species Clustering Efficiency

As illustrated in Figure 9b, classification performance was the lowest when utilizing context-independent embeddings generated by the pre-trained ESM-2. PST embeddings improved classification outcomes slightly, affirming that PST does capture genome-level context to some degree. However, embeddings enriched with extensive genomic context further enhanced classifier accuracy across all evaluated class sizes. Among the tested models, LV-3C consistently achieved superior performance, reflected in the highest F1-scores for every class configuration.

#### 4.4.3 Analysis of Embedding Variance and Distribution

The analysis (Figure 10) revealed distinct variability patterns among these embedding spaces. Embeddings produced by ESM-2 demonstrated the narrowest distribution, with the lowest interquartile range (1.16) and a relatively limited embedding space extent (range: 9.43). In contrast, embeddings from model LV-5B showed greater variability, with a broader interquartile range (1.58) and a larger embedding space (range: 15.69). Model LV-3C exhibited the most dispersed embeddings, spanning the widest embedding space (range: 16.18) and displaying significantly higher average pairwise Euclidean distances. These results indicate that incorporating genomic context into LV-3C substantially diversifies embedding representations, likely capturing biologically meaningful, genome-specific information.

**Figure 10.**
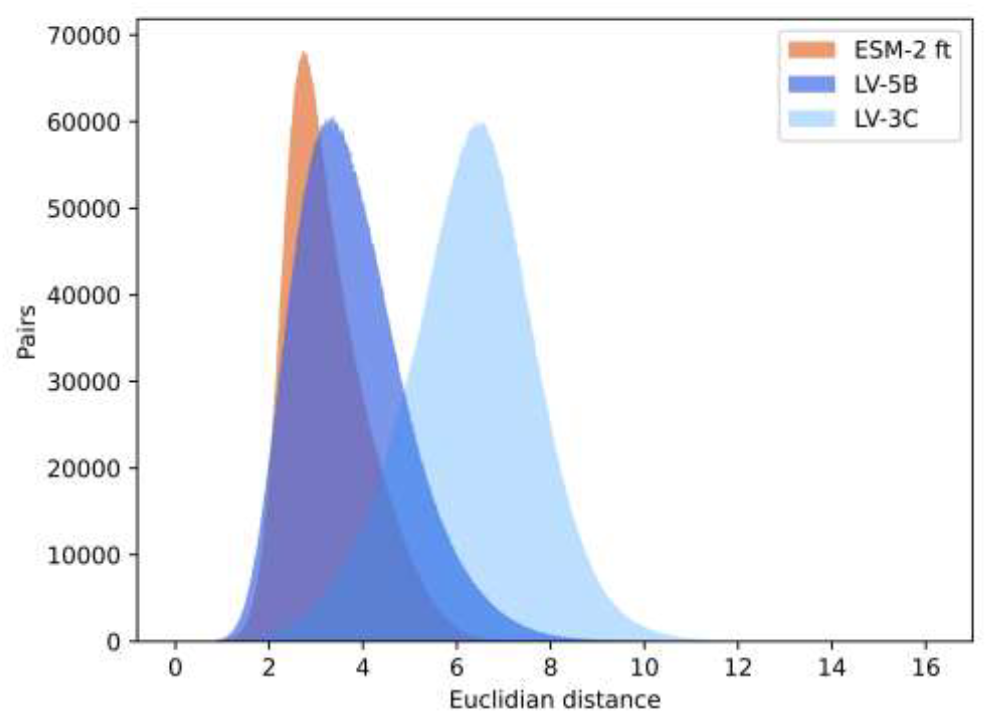
Protein Embeddings pairwise Euclidean distance. Each curve summarizes ≈12.5 million pair comparisons: lower distances (left tail) indicate within-species similarity, while greater distances reflect increasing taxonomic separation. ESM-2-ft yields the most compact manifold, LV-5B broadens the range, and LV-3C further widens and right-shifts the distribution, demonstrating its larger dynamic range for distinguishing distantly related proteins without sacrificing tight clustering of closely related sequences.

Further analysis of the Euclidean distance across protein embeddings shows that context provides a stronger correlation with taxonomic distance (Figures 9, 10). Specifically, embeddings from proteins within the same species exhibit high similarity, characterized by low Euclidean distance. As the taxonomic distance between proteins increases, embedding similarity correspondingly decreases, highlighting the models’ ability to capture biologically meaningful divergence across broader taxonomic categories.

### 4.5 Evaluating Across-Model Attention

The distribution of normalized attention scores across the three regions described in section 3.7 (short-, medium-and long-range) provides insight into the model’s capacity to capture biologically relevant interactions at different genomic scales. High variance in the attention score distribution indicates preferential focus on specific interaction ranges, while more uniform distributions suggest balanced attention across all distance categories. This analysis is particularly valuable for assessing whether the model effectively captures the long-range dependencies that conventional protein language models typically miss due to context limitations.

As Figure 11a shows, the attention profiles (short-, medium-, and long-distance) differ substantially among the four models evaluated. The pre-trained ESM-2 model exhibits a highly localized attention mechanism, concentrating almost exclusively on short-range interactions (100%), typically within individual proteins or immediate neighboring residues. This pattern aligns with ESM-2’s training paradigm, which focused on single-protein contexts and thus developed representations primarily capturing local residue relationships.

**Figure 11.**
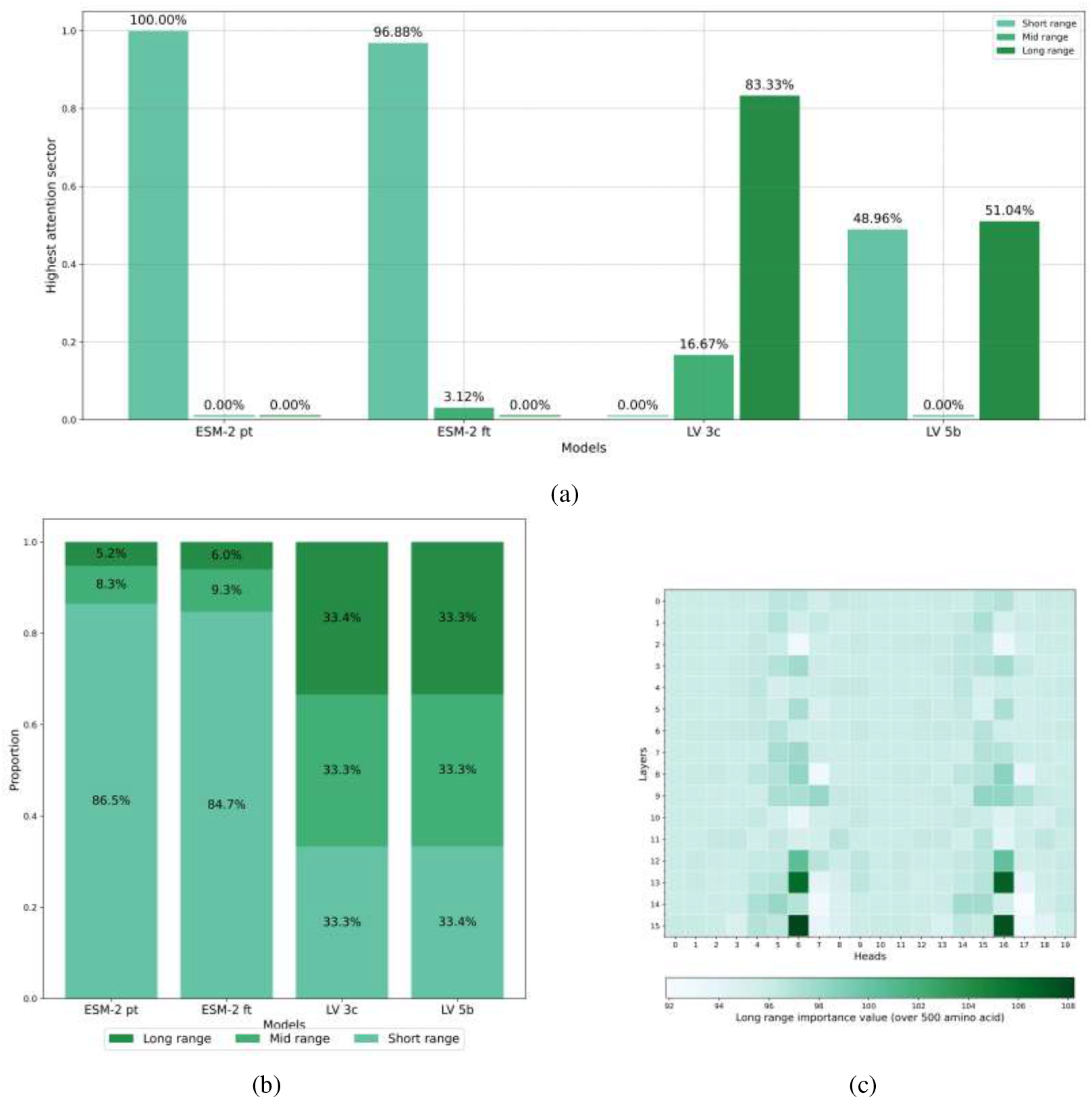
Attention analysis of our models. **(a)** Prevalence of dominant attention sectors (short-, medium-, or long-distance) across viral genomes. The chart represents 1,000 randomly selected viral genomes (≤ 13,000 aa in length), showing the proportion of genomes in which each interaction range category (short, medium, or long-distance) exhibited the highest mean attention score. **(b)** Distribution of normalized attention scores across interaction distance categories in viral proteomes. Stacked bar charts represent the distribution of attention scores for short-range (≤ 100 aa), medium-range (100–500 aa), and long-range (≥ 500 aa) interactions across 1,000 viral genomes. **(c)** Fine-grained analysis of long-range attention patterns in the LV-3C model. Darker (green) blocks indicate attention heads with a predominant long range attention. The visualization demonstrates how attention mechanisms capture biologically significant inter-protein interactions spanning substantial genomic distances within viral proteomes. Distribution variance indicates the relative focus of attention heads on specific interaction distances.

In contrast, the ESM-2 fine-tuned model, which employs a sliding-window (LongFormer) attention approach, demonstrates a modest expansion in attention scope. While it identifies a small fraction of medium-distance interactions (3.12%), it still predominantly focuses on short-distance interactions (96.88%). These findings suggest that although the sliding-window methodology enables detection of certain medium-range relationships, the model largely remains constrained to interactions between proximal amino acids.

Our models, LV-3C and LV-5B, exhibit markedly different attention distributions. Both models display balanced average attention scores across all distance categories (short, medium, and long), indicating a comprehensive approach to protein sequence modeling that equally integrates local and global contextual information. However, important distinctions between the two models emerge:

- Model LV-3C most frequently attributes maximal attention to long-distance interactions. This behavior highlights its enhanced capability to capture meaningful relationships between amino acids widely separated in sequence space, potentially spanning different proteins within the viral genome.
- Model LV-5B, while maintaining generally balanced average scores, frequently assigns highest attention importance to both short-and long-distance interactions. This indicates a flexible modeling strategy capable of emphasizing both local structural information and global protein–protein relationships as needed. Notably, LV-5B’s attention distribution retains traces of its ESM-2 initialization, suggesting it effectively combines characteristics of the pre-trained model with adaptations necessary for the expanded genomic context.

To further investigate inter-protein interactions, we calculated attention scores and identified attention heads specifically enriched for long-range information. Following the methodology described by [38], we systematically determined the protein pairs exhibiting the strongest interaction signals within viral genomes. To validate the biological relevance of these model-derived attention patterns, we benchmarked our predicted protein-protein interactions against experimentally validated interactions documented in the STRING database [45]. Our validation encompassed 146 annotated interactions across five distinct viral genomes.

In a binary classification framework, using a STRING interaction confidence threshold of 0.500, our attention-based method achieved an average F1 score of 0.74 and accuracy of 0.88. The classifier demonstrated an Area Under the Curve (AUC) of 0.66 (Figure 12, with a bootstrapped 95% confidence interval of [0.56, 0.75]), indicating statistically significant predictive performance. These results demonstrate substantial concordance between our attention-based predictions and experimentally validated interactions in the STRING database. The robust F1 score and high accuracy underscore our method’s capacity to identify biologically meaningful protein interactions within viral genomes. Furthermore, the moderate-to-strong positive correlation between attention scores and interaction confidence provides additional evidence for the biological relevance of the learned attention patterns.

**Figure 12.**
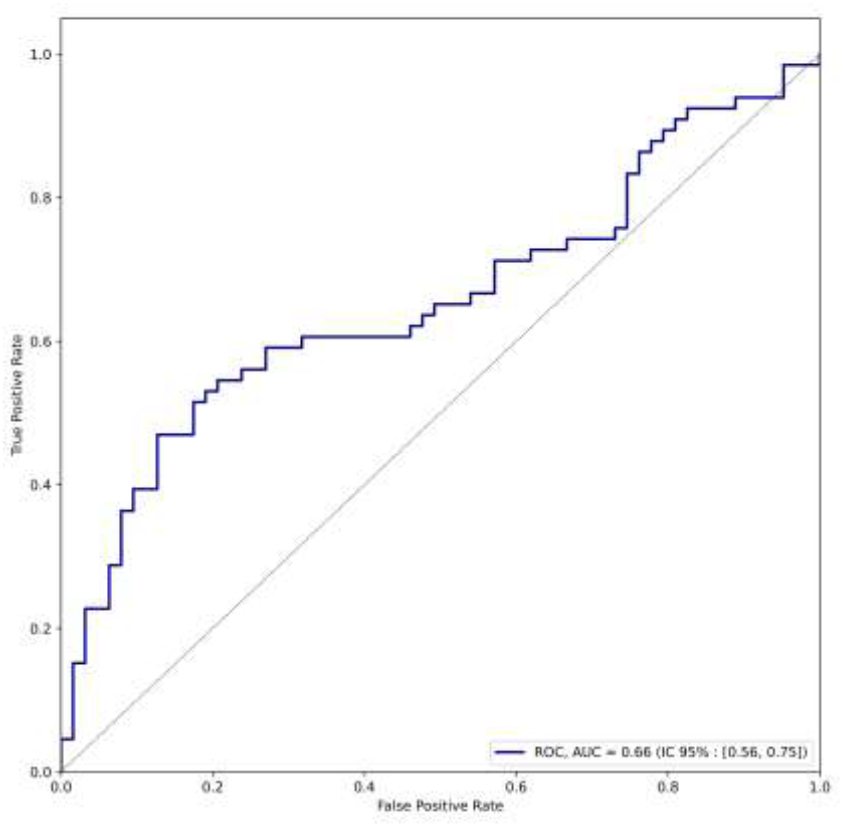
Receiver Operating Characteristic (ROC) curve of the binary classification task.

## 5 Discussion

Our findings demonstrate that extending transformer-based language models to encompass entire viral proteomes yields substantial improvements in both local amino acid prediction and broader protein function inference. By designing a content-aware sparse attention mechanism guided by inter-protein co-evolutionary signals, we have addressed the long-standing challenge of modeling densely packed viral genomes while maintaining computational tractability.

A central insight from our study is that higher-order genomic context—the collective presence and arrangement of multiple proteins—significantly enhances protein representation accuracy beyond what can be achieved by analyzing protein sequences in isolation [3], or using context length extension [15, 16] or context-agnostic sparse attention [25, 26, 27]. As the results show, conventional protein language models trained on individual proteins (e.g., ESM-2) focus on local sequence features and exhibit marked performance degradation when context extends beyond their trained window. In contrast, our biologically informed sparse attention approach captures both local relationships and long-range dependencies across entire viral genomes. This more comprehensive perspective reveals functional couplings between non-consecutive proteins that would otherwise remain undetected, such as the well-documented interactions between DNA polymerase I, ribonucleotide reductase, and helicase proteins [41, 42, 43].

The attention patterns learned by our models demonstrate remarkable alignment with experimentally validated protein-protein interactions, benchmarked against the STRING database [45]. This suggests that attention mechanisms in our models effectively capture biologically meaningful co-evolutionary relationships, providing a computational approach to predict functional protein interactions that complements experimental techniques. The balanced distribution of attention across short, medium, and long-range interactions in our LV-3C and LV-5B models – in contrast to the predominantly local focus to other models – further reinforces the biological relevance of our approach.

The significant divergence in embedding spaces between context-independent and context-aware models, as evidenced by the distinct distributions of pairwise Euclidean distances, underscores the fundamental difference in information captured by these approaches. The wider embedding space of our LV-3C model, with its greater interquartile range and broader distribution of distances, suggests a richer representation that incorporates both sequence-specific and context-dependent information.

Despite these advances, computational overhead remains a practical consideration. Our selection of 50 protein-protein interactions for the sparse attention mechanism strikes an effective balance for viral datasets while maintaining computational efficiency. This parameter can be readily increased if GPU resources are available. We intentionally constrained our implementation to run on a single GPU, as we believe this represents a widely accessible computational environment for most research groups. While we utilized an NVIDIA A6000 for this study, the rapid advancement of more affordable GPUs (sub-$5,000) with comparable memory capacities makes our approach increasingly accessible. For processing significantly larger genomes—particularly from eukaryotic organisms—additional optimizations may be beneficial. Additionally, our semi-supervised approach to inferring putative interactions could benefit from larger ground-truth databases or complementary evidence from proteomic or structural studies to further refine the biological relevance of the learned attention patterns.

Looking forward, our framework offers considerable promise for extension to larger viral genomes, bacteriophages, and potentially even eukaryotic genomes with appropriate scaling. The methodology could be particularly valuable for analyzing metagenomic data, where contextual information might help resolve ambiguities in protein function assignment. Furthermore, the correlation between embedding similarity and taxonomic distance suggests potential applications in phylogenetic analysis and evolutionary studies.

## 6 Conclusion

Our study proposes a novel framework for viral protein language modeling by extending transformer architectures to process entire viral genomes. By integrating a biologically informed sparse attention mechanism, our approach efficiently captures long-range inter-protein interactions that are missed by conventional single-protein models. This design not only reduces perplexity across extended sequences (up to 61,000 aa) but also generates context-aware embeddings that accurately recapitulate known co-evolutionary relationships—for instance, among PolA, RNR, and helicase.

Our results demonstrate that large-scale learning improves embedding quality. The enhanced clustering of protein and genome embeddings, along with robust taxonomic classification, underscores the broader applicability of our method to phylogenetic analysis and metagenomic studies. Our work provides a solid foundation working with viral genomes and paves the way for more refined analyses of protein–protein interactions, ultimately deepening our understanding of viral biology and evolution.

Looking ahead, our framework offers considerable promise for extension to larger viral genomes, bacteriophages, and potentially even eukaryotic genomes. By refining the biological specificity of sparse attention patterns and incorporating additional layers of experimental validation, our approach can continue to evolve toward more precise and comprehensive genomic analyses. Ultimately, this work not only deepens our understanding of viral genome organization and evolution but also provides a scalable framework for protein modeling at genomic scales, bridging the gap between computational biology and experimental validation.

## Supporting information

Supplementary Material: NCBI Experiment

## 7 Acknowledgments

This work is supported by the National Science Foundation grants 1736030 and 2025567. Support from the University of Delaware Center for Bioinformatics and Computational Biology (CBCB) Core Facility, the University of Delaware Sequencing and Genotyping Center, and use of the BIOMIX compute cluster was made possible through funding from Delaware INBRE (NIGMS P20GM103446, NIH S10OD028725), and the State of Delaware.

