## Supplementary Material: NCBI Experiment for "Extending Protein Language Models to a Viral Genomic Scale Using Biologically Induced Sparse Attention"

### Using embeddings to predict taxonomy

To assess the taxonomic discriminability of the learned embeddings, we conducted a supervised classification analysis. For each protein or genome in the dataset, a 1,280-dimensional embedding vector was generated and paired with its corresponding taxonomic label. The dataset, comprising 100,000 proteins and 5,000 genomes, was randomly partitioned into training (80%) and validation (20%) subsets. To ensure a robust evaluation, we restricted the analysis to the top X most frequently represented taxonomic species. A single-layer neural network with an intermediate hidden dimension of 2,560 was trained for 1,000 epochs using a learning rate of 1e-3. Model performance was evaluated using 5-fold cross-validation, and standard classification metrics were reported.

### NCBI Virus Database

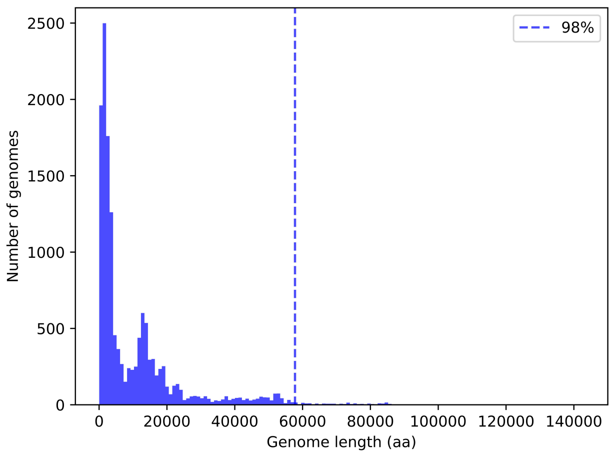


Figure 13: **Viral genome length distributions in the NCBI Virus Database.** This histogram shows that a context length of 61k amino acid allows us to process over 98% of the 14,436 complete geomes of the database.
